# Proteomic Profiling of Mesenchymal Stem Cell-Derived Extracellular Vesicles: Impact of Isolation Methods on Protein Cargo

**DOI:** 10.1101/2024.05.06.592788

**Authors:** Morteza Abyadeh, Shahab Mirshahvaladi, Sara Assar Kashani, Joao A. Paulo, Ardeshir Amirkhani, Fatemeh Mehryab, Homeira Seidi, Niloufar Moradpour, Sheyda Jodeiryjabarzade, Mehdi Mirzaei, Vivek Gupta, Faezeh Shekari, Ghasem Hosseini Salekdeh

**Author notes:** Corresponding authors: Faezeh Shekari, Department of Stem Cells and Developmental Biology, Cell Science ResearchCenter, Royan Institute for Stem Cell Biology and Technology, ACECR, Tehran, Iran; Ghasem Hosseini Salekdeh, Faculty of Natural Sciences, Macquarie University, North Ryde, NSW, Australia.

## Abstract

Extracellular vesicles (EVs) are nanosized vesicles with a lipid bilayer that are secreted by cells and play a critical role in cell-to-cell communication. Despite the promising reports regarding their diagnostic and therapeutic potential, the utilization of EVs in the clinical setting is limited due to insufficient information about their cargo and a lack of standardization in isolation and analysis methods. Considering protein cargos in EVs as key contributors to their therapeutic potency, we conducted a tandem mass tag (TMT) quantitative proteomics analysis of three subpopulations of mesenchymal stem cell (MSC)-derived EVs obtained through three different isolation techniques: ultracentrifugation (UC), high-speed centrifugation (HS), and ultracentrifugation on sucrose cushion (SU). Subsequently, we checked EV marker expression, size distribution, and morphological characterization, followed by bioinformatic analysis. The bioinformatic analysis of the proteome results revealed that these subpopulations exhibit distinct molecular and functional characteristics. The choice of isolation method impacts the proteome of isolated EVs by isolating different subpopulations of EVs. Specifically, EVs isolated through the high-speed centrifugation (HS) method exhibited a higher abundance of ribosomal and mitochondrial proteins. Functional apoptosis assays comparing isolated mitochondria with different EV isolation methods revealed that HS-EVs, but not other EVs, induced early apoptosis in cancer cells. On the other hand, EVs isolated using the sucrose cushion (SU) and ultracentrifugation (UC) methods demonstrated a higher abundance of proteins primarily involved in the immune response, cellLJcell interactions, and extracellular matrix interactions. Our analyses unveil notable disparities in proteins and associated biological functions among EV subpopulations, underscoring the importance of meticulously selecting isolation methods and resultant EV subpopulations based on the intended application.

## 1 INTRODUCTION

Extracellular vesicles (EVs) are small membrane-enclosed vesicles secreted by most cell types that play a crucial role in intercellular communication (Cheng & Hill, 2022; M. Park et al., 2021). EVs contain various types of biomolecules, including proteins, nucleic acids, and lipids, that can be transferred to other cells and modulate their physiological functions (Murphy et al., 2019; Faezeh Shekari, Morteza Abyadeh, et al., 2023). EVs have been shown to be involved in various physiological and pathological processes, such as immune response, cancer progression, and neurodegeneration (Beretta et al., 2020; Gupta et al., 2022; Gurung, Perocheau, Touramanidou, & Baruteau, 2021; M. Park et al., 2021; Ruan et al., 2021). These lipid-bound particles have also emerged as promising candidates for therapeutics due to their unique properties, such as biocompatibility, low immunogenicity, and the ability to cross biological barriers (Abyadeh et al., 2023; Barile et al., 2014; Haghighitalab et al., 2023). They can also target specific cells and tissues, making them an attractive alternative to liposomes (Herrmann, Wood, & Fuhrmann, 2021; Mehryab et al., 2020). According to a scientific consensus, EVs are classified into three subtypes based on their biogenesis. These incude exosomes, ectosomes and apoptotic bodies that derived from endosomal compartments, plasma membrane budding, and dying cells shedding respectively (Théry et al., 2018; Witwer & Thery, 2019).

A growing number of studies have reported on the diagnostic and therapeutic potential of EVs in various devastating diseases, including cancer, cardiovascular diseases, and neurodegenerative disorders. Given the diverse nature of EVs, the variability in administration routes, and the differences in disease models used, the published results lack direct comparability (F. Shekari et al., 2023; F. Shekari et al., 2021; Zhang et al., 2022). Therefore, this report aimed to uncover some aspects of EV heterogeneity by analyzing their proteome profiles.

Disease diagnosis and treatment with EVs is a new and emerging field that faces several challenges. Specifically, the main challenge currently is improving and standardizing methods for EV isolation and analysis to use them effectively for therapeutic purposes (Dash, Palaniyandi, Ramalingam, Sahabudeen, & Raja, 2021; Li, Kaslan, Lee, Yao, & Gao, 2017; F. Shekari et al., 2023; Shirejini & Inci, 2022; Singh et al., 2021). Isolation of EVs from biological fluids or cell cultures is a critical step for their characterization and functional studies. Currently, methods commonly utilized to isolate EVs include ultracentrifugation, size-exclusion chromatography, and precipitation-based methods (Y. B. Kim, Lee, & Moon, 2022; Li et al., 2017; Faezeh Shekari, Morteza Abyadeh, et al., 2023). The choice of isolation method for EVs can have a significant impact on the quality and quantity of the obtained EVs as well as their cargo composition (Poupardin et al., 2024). EVs serve as vehicles for the transfer of a diverse cargo, both internally and on their surface including a wide array of biomolecules, including proteins, lipids, sugars, small molecules, and various nucleic acid species (Dixson, Dawson, Di Vizio, & Weaver, 2023; Tóth et al., 2021). Using different methods results in EVs with different features, such as different sizes, heterogeneity and morphology, that may affect their content and subsequently their biological characteristics (Dash et al., 2021; Konoshenko, Lekchnov, Vlassov, & Laktionov, 2018; Yamashita, Takahashi, Nishikawa, & Takakura, 2016). Recent studies have highlighted that EVs isolated from human monocytic cell lines, plasma, and cancer cells through various isolation methods exhibit variations in sizes and distinct protein contents. This variation in protein content can potentially impact the therapeutic properties. These differences can serve as markers for subpopulations of EVs (Jimenez et al., 2019; Lischnig, Bergqvist, Ochiya, & Lässer, 2022; Veerman et al., 2021). These studies used different cancer cell lines, different centrifuge speeds (including 10,000 × g and 16,500× g for large vesicle isolation and 100,000 × g and 118,000 × g for small vesicle isolation), and employed different definitions for small and large vesicles. Despite the variations in published results so far they generally indicate that large EVs tend to contain more ribosomal proteins and those involved in translation initiation and metabolism, whereas small EVs are enriched with proteins related to cellLJcell adhesion and extracellular matrix organization. (Jimenez et al., 2019; Lischnig et al., 2022). In this study, we performed a proteomic analysis of mesenchymal stem cell (MSC)-derived EVs isolated using different isolation methods, including high-speed centrifugation (HS), ultracentrifugation (UC), and ultracentrifugation on sucrose cushion (SU), to investigate the effect of isolation method on the proteome of EVs.

MSCs are well known for their anti-inflammatory effects. MSCs interact with immune cells in both the innate and adaptive immune systems, making them a potential therapy against many diseases, and nearly 60 clinical trials have been approved by the U.S. FDA (Kou et al., 2022; Wiest & Zubair, 2020; Xiaomo Wu et al., 2020). A wealth of evidence indicates that the therapeutic effects of MSCs are attributed to the paracrine-like secretion of cytokines (including growth factors and chemokines) and EVs with their involvement in cellular communication (Arabpour, Saghazadeh, & Rezaei, 2021; Asgarpour et al., 2020; Furuta et al., 2016; Rahmani et al., 2020). It has also been reported that EVs derived from MSCs retain the therapeutic properties of the parent MSCs (F. Shekari et al., 2021). Our previous researches have highlighted the therapeutic efficacy of UC-EVs in chronic liver failure, showcasing significant antifibrotic, anti-inflammatory, anti-apoptotic, and regenerative effects (Mardpour et al., 2019). In a mouse model of optic nerve crush, treatment with UC-EVs led to notable improvements in GAP43+ axon counts and cognitive visual behavior (Seyedrazizadeh et al., 2020). Furthermore, SU EVs exhibited effectiveness in mitigating cistauosis and apoptosis, enhancing outcomes in a mouse model of weight drop-induced traumatic brain injury (Amini et al., 2023), and mitigate clinical symptoms in non-human primates with inflammatory-mediated type 1 diabetes through immunomodulation (Kashani et al., 2023). When administered via intramyocardial injection in rats with myocardial infarction, UC-EVs prevented severe ventricular remodeling (Firoozi et al., 2020). Recognizing the challenges associated with isolating UC EVs for clinical trials, particularly regarding the feasibility and cost-effectiveness of adhering to GMP standards, we studied HS EVs. These HS-EVs showed efficacy in improving ovarian function and restoring fertility in a mouse model of chemotherapy-induced premature ovarian failure (Eslami et al., 2023). Additionally, HS EVs successfully restored endometrial function in an experimental rat model of Asherman syndrome (Mansouri-Kivaj et al., 2023). In a phase 1 clinical trial, the combined administration of MSCs and HS-MSC-EVs significantly reduced serum levels of pro-inflammatory markers in COVID-19 patients, with no reported serious adverse events (Zarrabi et al., 2023). Herein, our results revealed distinct proteome profiles within the EVs based on various centrifugation-based isolation methods. Specifically, EVs isolated through the HS method exhibited a higher abundance of proteins involved in energy metabolism-related pathways, regulation of the actin cytoskeleton, tight junctions, and axon guidance than EVs isolated using the SU and UC methods. Conversely, EVs isolated through the SU or UC methods showed a higher abundance of proteins involved in inflammatory responses, complement and coagulation cascades, Th17 cell differentiation, neutrophil extracellular traps, and cell death. These findings suggest that the choice of isolation method can influence not only the quantity and quality of EVs but also their cargo composition. Our study highlights the importance of carefully selecting EV isolation methods for future downstream applications.

## 2 MATERIALS AND METHODS

### 2.1 Culture and characterization of clonal mesenchymal stromal cells (MSCs)

Human bone marrow clonal MSCs were provided by the Royan Stem Cell Bank (RSCB0178), Tehran, Iran. cMSCs were established based on the subfractionation culturing method (Song et al., 2008; Yi et al., 2015). In this investigation, a two-dimensional (2D) cell culture approach was employed. In brief, single colony-forming unit-derived colonies were produced by culturing bone marrow in culture dishes containing low-glucose Dulbecco’s modified Eagle’s medium (DMEM-LG; 10567022, Thermo Fisher) supplemented with 20% fetal bovine serum (FBS; SH30071.03IR, HyClone) and 2 mM GlutaMAX (Life Technologies) in a 37 °C and 5% CO2 incubator. To increase the homogeneity of the MSC products, the eligible CFU were chosen for culture. The MSCs were characterized at passage ten by an MSC Phenotyping Kit (130-095-198, MACS Miltenyi Biotec). The kit contained anti-CD73, CD90, CD105 CD14, CD20, CD34, and CD45 surface markers and was analyzed by flow cytometry (BD FACSCalibur™). Each surface marker percentage was calculated with Flowing software version 2.5.1. MSCs were also evaluated by lineage differentiation into osteocytes (A1007201, Thermo Fisher), adipocytes (A1007001, Thermo Fisher) and chondrocytes (A1007101, Thermo Fisher).

### 2.2 Isolation of mesenchymal stromal cell-derived extracellular vesicles (MSC-EVs)

EVs were isolated by three different methods, including HS, UC, and SU as shown in Fig 1A. Briefly, cMSCs were cultured to 80%-90% confluence in a T75 culture flask and after 48 hrs approximately 500 ml MSC-conditioned media (CM) corresponding to 75 × 106 cells were collected and frozen at −80 °C for EV isolation. The MSC-CM was centrifuged at 3,000 × g for 10 min to remove debris and dead cells and then at 20,000 × g to pellet the HS fraction. The collected CM from the previous step was subjected to UC and SU centrifugation to isolate UC and SU EVs, respectively. For the UC isolation method, half of the supernatant was subjected to 110,000 × g centrifugation (Beckman Coulter, Kraemer Boulevard Brea, CA) for 2 hrs at 4 °C. The pellet of this step was washed with phosphate-buffered saline (PBS) (10010049, Gibco) using UC for an additional 2 hrs. The other half of the CM was subjected to a 65% sucrose gradient at 110,000 × g for 2 hrs. The culminated interface was washed with PBS for an additional 2 hrs at 110,000 × g. The final products were stored at −80 °C for future use.

**Figure 1.**
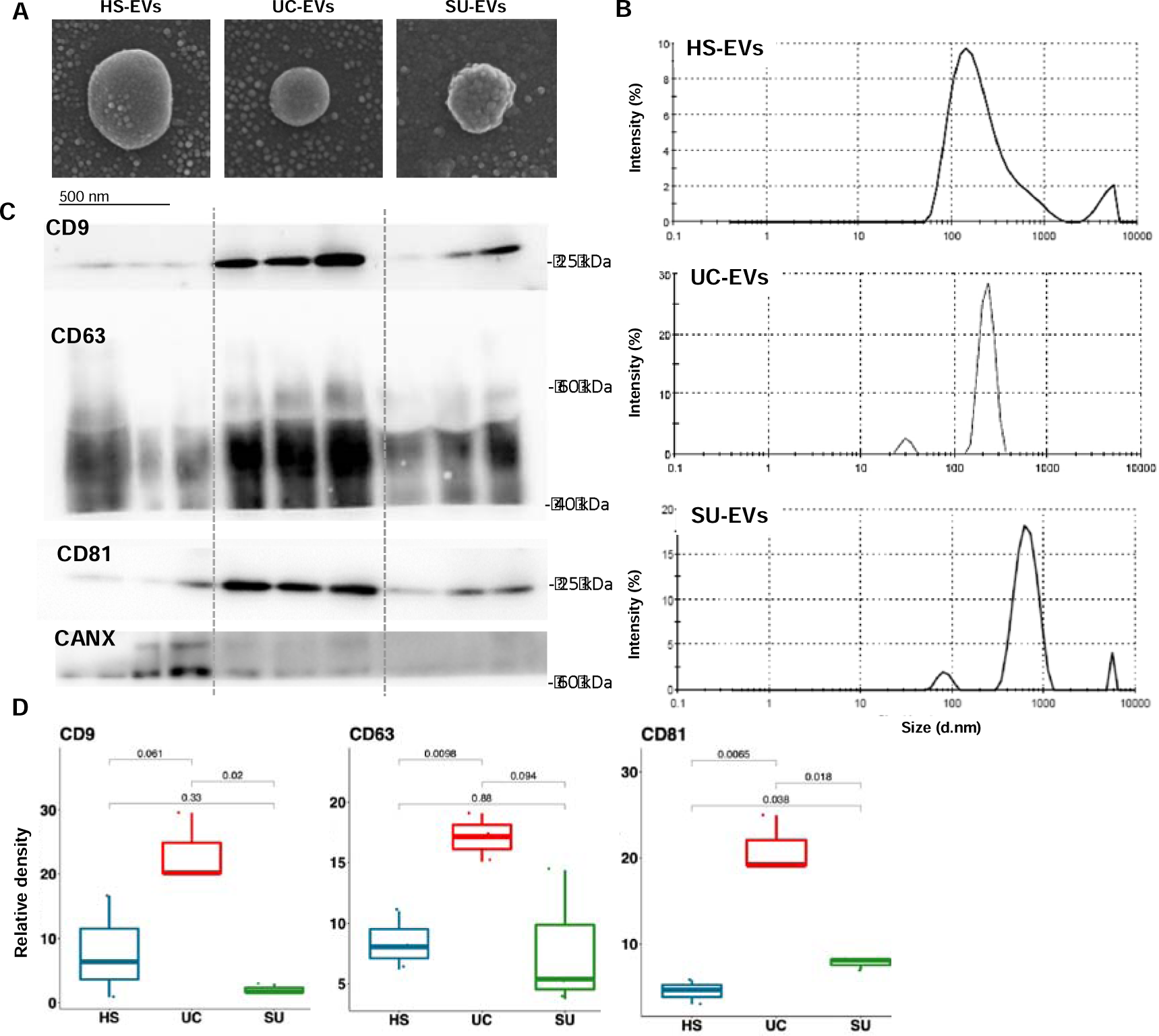
Flowchart of isolation (A) and characterization of cMSC-derived extracellular vesicles (cMSC-EVs). (B) Morphology assessment of EVs at the single-vesicle level by scanning electron microscopy, scale bar: 500 nm. (C) Western blotting analysis for enriched EV markers for each subpopulation for CD9, CD63, CD81, and calnexin (CANX) as negative markers in three replicates. (D) Size distribution assessment of EVs by dynamic light scattering. (E) The box chart shows the quantification of protein bands, and the results are expressed as the middle line at the mean. P values were calculated by the t test in the R platform. HS: EVs isolated by 20 Kg, UC: EVs isolated by ultracentrifugation, SU: EVs isolated by ultracentrifugation on sucrose gradient

### 2.3 Western blotting

Protein expression of selected EV markers was evaluated using WB analysis as described previously(Tahamtani et al., 2014). The protein concentration of each sample was determined using a BCA assay kit (Pierce) and a MULTISKAN microplate reader (Thermo Scientific). Samples (20-25 μg) were separated by SDSLJPAGE electrophoresis (10%). The gels were run according to standard methods, and the probed proteins were transferred to Bio-Rad polyvinylidene fluoride membranes using a semidry electroblotting transfer system (Bio-Rad) at 25 V for 2 h and 30 min at room temperature. After the PVDF membranes were blocked with 2% BSA in Tris-buffered saline plus 0.1% Tween for 1 h, then were incubated overnight at 4 °C with the following primary antibodies: anti-CD9 (Sc13118, Santa Cruz; 1:1000), anti-CD81 (Sc7637, Santa Cruz; 1:1000), anti-CD63 (Sc-5275, Santa Cruz; 1:200), anti-TSG101 (Gtx70255, Genetex; 1:1000) and anti-calnexin (Sc11397, Santa Cruz; 1:1000) as a negative marker. The membranes were washed three times with TBST and then incubated with horseradish peroxidase (HRP)-conjugated secondary antibodies (diluted at 1:100,000) at room temperature for 1 h. After three TBST rinses, the membranes were incubated with SuperSignalWest Femto Substrate (Thermo Fisher Scientific), and bands were visualized using the Alliance Q9 Advanced Chemiluminescence Imager gel documentation system. The band intensities were normalized to the loading control and quantified using ImageJ (Schneider, Rasband, & Eliceiri, 2012) software version 1.47. The quantification was conducted in triplicate to serve as analytical replication.

### 2.4 Scanning Electron Microscopy (SEM)

Each MSC-EV pellet was vortexed and resuspended in 0.2 ml of PBS. From each sample, 1-5 µl droplets were immobilized on clean slides and blown dry under a fume hood. The samples were imaged within 2 days by scanning electron microscopy (SEM Model QUANTA 200, FEI, USA). Then, a thin layer of gold was applied to the sample using a specialized coating device (Nanostructured Coatings Co., Iran). The samples were imaged by scanning electron microscopy (MIRA3 (TESCAN, Czech Republic).

### 2.5 Dynamic light scattering (DLS)

To determine the size of MSC-EV subpopulations, we used DLS with an SZ-100z Dynamic Light Scattering & Zeta potential analyzer (Horiba Jobin Jyovin, λ: 532 nm laser wavelength). Five µg of each subpopulation was resuspended in 1 ml of PBS in a removable clean cuvette. The size reported is based on the intensity.

### 2.6 Enzyme-linked immunosorbent assay (ELISA)

The quantity of EV protein was measured using a BCA assay, and 20 µg of each EV subpopulation was analyzed for the presence of human insulin-like growth Factor 1 (IGF-1; DG100B, R & D systems) according to the manufacturer’s protocol. Briefly, samples were incubated with 50 μl of assay diluent in ELISA plates for 3 h at 2-8 °C. Following three washes with 400 μl washing buffer each (up to 2 min/each), the plate was subjected to one-hour human IGF-1 conjugate at room temperature. After washing as explained before, the subsequent steps, including substrate addition, enzymatic reaction termination, and absorbance measurement at 450 nm, were performed according to the manufacturer’s instructions. All EV samples were analyzed in triplicate.

### 2.7 Coagulation assays

Blood samples were collected using vacuum blood collection tubes (Vitrex, Denmark), with each tube containing 2.7 mL of blood. Subsequently, 100 µg of EV sample was added to each tube, while another group was treated with PBS as a control. Within an hour, the fresh blood was centrifuged at 3000 rpm for 10 minutes to obtain plasma. Prothrombin time (PT) and activated partial thromboplastin time (aPTT) were analyzed at the local laboratory of Royan hospital (Royan Institute, Iran).

### 2.8 Lymphocyte proliferation assay (LPA)

Blood samples were collected using vacuum blood collection tubes (Vitrex, Denmark), and peripheral blood mononuclear cells (PBMCs) were extracted from healthy donors’ buffy coats. Carboxyfluorescein succinimidyl ester (CFSE; Invitrogen, USA) pre-labeled T lymphocytes were then cultured in a 96 round-bottom well plate (1.5×105 cells per well) in supplemented RPMI 1640 medium (Gibco, UK) and stimulated with phytohemagglutinin (PHA; Sigma-Aldrich, USA). The lymphocytes were incubated for 4 days and treated with 10 µg of EV samples on the 1st and 3rd days. Live cells were identified by propidium iodide (PI; Sigma-Aldrich, USA) staining, and cell proliferation was assessed by visualizing cell generations using CFSE flow cytometry on a BD FACSCalibur (BD Biosciences, USA).

### 2.9 NanoLCDMS/MS and tandem mass tag (TMT) quantitative proteomics

The protein was processed for mass spectrometric analysis using commercially procured S-Traps (Protifi, USA) according to the manufacturer’s instructions. Briefly, protein powder was resuspended in 1× SDS lysis buffer (5% SDS, 100 mM triethylammonium bicarbonate, TEAB, pH 7.55), and the concentration was measured using a BCA assay (Thermo Scientific, USA). Equal amounts (50 µg) of samples were aliquoted, and volumes were normalized across all the samples using S-Trap lysis buffer (5% SDS, 100 mM triethylammonium bicarbonate, TEAB, pH 7.55). To enable dissolution of the protein, pellets were vortexed well. Cysteine disulfide bonds of the proteins were reduced with 10 mM DTT at 95 °C for 10 min and then alkylated with 25 mM IAA for 30 min in the dark at room temperature. The pH of the samples was adjusted using 12% aqueous phosphoric acid, added at 1:10 for a final concentration of ∼1.2% phosphoric acid and diluted using S-Trap binding buffer (90% aqueous methanol containing a final concentration of 100 mM TEAB, pH 7.55). S-Trap binding buffer was added to the acidified lysis buffer, and the sample mixture was transferred to a labeled S-Trap column and centrifuged at 4,000 g, after which the flow through was discarded. The column was washed twice using S-Trap binding buffer, and proteins retained on the column were digested in the presence of 20 µL trypsin solution (1:25 trypsin to protein ratio, total ∼2 µg trypsin in 50 mM triethylammonium bicarbonate) for 1 hour at 47 °C. Following digestion, peptides were eluted off the column after the addition of 50 mM triethylammonium bicarbonate and centrifugation. The remaining peptides were eluted from the column using sequential centrifugation with the addition of 0.2% aqueous formic acid followed by 50% aqueous ACN containing 0.2% formic acid. Peptides were dried by vacuum centrifugation and then reconstituted in 0.1% formic acid. The peptide concentration was determined using the Pierce quantitative colorimetric peptide assay (Thermo Scientific, USA).

TMT reagent labeling of peptides was performed according to the manufacturer’s (Thermo Scientific, USA) instructions. Briefly, anhydrous acetonitrile was added to each TMT-labeled vial, followed by vortexing for 5 min and brief centrifugation. Aliquots of individual peptide samples were labeled with each of the individual TMT labels (total of ten labels). Labeling was performed at room temperature for 1 h with occasional vortexing. To quench the excess TMT label in the sample, 5% hydroxylamine was added to each of the samples, vortexed, and then incubated at room temperature for 15 min. Before pooling the samples and to ensure that equal amounts of total peptides were pooled from all samples, a “label check” experiment was performed by mixing 1.5 μL of each individually labeled TMT sample. The “label check” sample was vacuum dried and then desalted using a solid phase extraction disk Styrene Divinyl Benzene containing Stage tips (Empore SDB-RPS 47 mm extraction disk, SUPLCO). Samples were reconstituted in 2% ACN and 0.1% FA in water after vacuum drying and analyzed by LC coupled to a mass spectrometer (Q-Exactive, Thermo Fisher, USA). A normalization factor was obtained from the label check experiment, and the TMT-labeled peptide samples were pooled at a 1:1 ratio across all samples and vacuum dried. The samples underwent purification through desalting using C18 solid-phase extraction (SPE) columns (Sep-Pak, Waters) and were then dried completely using vacuum centrifugation. The peptide mixture was fractionated by high pH (HpH) reversed-phase HPLC into 96 fractions that were then consolidated into 17 fractions prior to LCLJMS/MS analysis.

Cleaned peptides from each fraction were analyzed using a Q Exactive Orbitrap mass spectrometer (MS; Thermo Scientific) coupled to an EASY-nLC1000 nanoflow HPLC system (Thermo Scientific). Reversed-phase chromatographic separation was performed on an in-house packed reverse-phase column (75 μmLJ×LJ10 cm Halo 2.7-μm 160 Å ES-C18, Advanced Materials Technology). Labeled peptides were separated for 2 h using a gradient of 1%–30% solvent B (99.9% acetonitrile/0.1% formic acid) and Solvent A (97.9% water/2% acetonitrile/0.1% formic acid). The Q Exactive MS was operated in the data-dependent acquisition mode to automatically switch between full MS and MS/MS acquisition. Following the full MS scan from m/z 350–1850, MS/MS spectra were acquired at a resolution of 70,000 at m/z 400 and an automatic gain control target value of 106 ions. The top ten most abundant ions were selected with a precursor isolation width of 0.7 m/z for higher energy collisional dissociation (HCD) fragmentation. HCD-normalized collision energy was set to 35%, and fragmentation ions were detected in the Orbitrap at a resolution of 70,000. Target ions that had been selected for MS/MS were dynamically excluded for 90 s.

### 2.10 Proteomics data analysis

The mass spectrometric data files were searched using Proteome Discoverer (version 2.1, Thermo Scientific). The data were processed using the search engines SequestHT and Mascot (Matrix Science, London, UK) against human sequences. The quantitative ratios were generated using the raw abundance values in each channel. The MS1 tolerance was set toLJ±LJ10 ppm and the MS/MS tolerance to 0.02 Da, and trypsin digestion was specified with one missed cleavage allowed. Carbamidomethylation of cysteine and 10-plex TMT tags on lysine residues and peptide N-termini were set as static modifications. Oxidation of methionine and deamination of asparagine and glutamine residues were set as variable modifications. Search result filters were selected as follows: only peptides with a Mascot score > 15 and below the significance threshold filter of p = 0.05 were included, and single peptide identifications required a score equal to or above the Mascot identity threshold. The false discovery rate was set to 0.01 or less in Proteome Discoverer, and protein grouping was enabled such that when a set of peptides in one protein were equal to, or wholly contained within, the set of peptides of another protein, the two proteins were contained together in one protein group. Relative quantitation of peptides and proteins was achieved by pairwise comparison of TMT reporter ion intensities after normalization to the pooled internal standard.

TMTPrepPro (Mirzaei et al., 2017) was used for further analysis of the identified proteins. All protein ratios relative to the reference (label-131) were extracted. Proteins were considered differentially abundant if log2FC>|0.263| and p value < 0.05.

### 2.11 Functional enrichment analysis

For functional enrichment analysis of differentially abundant proteins (DAPs) in EV groups, we utilized Enrichr, a web-based tool for comprehensive gene sets (E. Y. Chen et al., 2013). KEGG (Kyoto Encyclopedia of Genes and Genomes) pathway, Gene Ontology (GO) biological process (BP), GO molecular function (MF), and GO cellular component (CC) analyses were then used to find the enriched terms from the submitted list of DAPs. Enriched terms with an adjusted p value less than 0.05 were considered statistically significant for DAPs.

### 2.12 Protein**D**protein interaction (PPI) network and hub gene analyses

ProteinLJprotein interaction (PPI) networks were analyzed using the Cytoscape-String App plugin with a confidence score > 0.5, as previously described (Abyadeh et al., 2022). Briefly, DAPs from each group were uploaded into Cytoscape. Next, the Homo sapiens database in StringDB was selected to generate the protein interactions between DAPs. To identify the hub genes within the protein network, CytoHubbaplugin in Cytoscape was utilized, and hub genes were selected based on the maximal clique centrality (MCC) algorithm (Chin et al., 2014).

### 2.13 Isolation of mitochondria from MSCs

MSCs was obtained from the Royan cell bank. Following de-freezing, cells were centrifuged at 1500 g for 5 minutes, and washed with PBS. Sucrose buffer was prepared (0.42 g sucrose and 0.05 g HEPES, 40 µl protease inhibitor in 5 ml PBS) and filtered through the 0.22 filter. Cell pellet was vortexed for 5 seconds and rested for 5 seconds until the cell suspension was lysed. Then sonication was performed at medium speed for 7 minutes with 30 seconds on and 30 seconds off. To ensure the cells were lysed, we observed them under an inverted microscope and compared them with the intact cells we had previously taken. After sonication, centrifugation steps have been performed based on our previously reported protocol (Faezeh Shekari et al., 2017; Tasbihi, Shekari, Hajjaran, Khanmohammadi, & Hadighi, 2020; Tasbihi, Shekari, Hajjaran, Masoori, & Hadighi, 2019). Briefly, the homogenate suspension of sonicated cells was centrifuged at 4°C for 10 minutes at 800 x g to separate any large particles including not broken cells or aggregates. Subsequently, the supernatant was further centrifuged for 15 minutes at 3000 x g to pellet smaller debris. Finally, the resulting supernatant underwent a final centrifugation step of 25 minutes at 18000 x g to isolate mitochondria. The mitochondria containing pellet is stored at −80 °C for future use.

### 2.14 Treatment with GW4869

A total of 1 × 10^6^ cMSCs were initially cultured in T75 cell culture flasks. Upon reaching 80% confluence, the complete medium of cMSCs was replaced with a medium containing 10µM GW4869 (CAS 6823-69-4), an inhibitor of exosome biogenesis/release (J. H. Kim, Lee, & Baek, 2022; Raiter, Lipovetsky, Stenbac, Lubin, & Yerushalmi, 2023). After 24 hrs, when the cells in the control group had reached 100% confluence, the culture media from both the control and GW4869-treated groups were collected. Subsequently, EVs were isolated from these media using two distinct methods: ultracentrifugation and high-speed centrifugation.

### 2.15 Meta-analysis and mitochondrial protein prediction

To validate our findings regarding the variable presence of mitochondrial proteins in distinct subpopulations of EVs, we conducted a comprehensive meta-analysis of previously reported proteomes. Supplementary Table 1 presents the proteome profiles of MSC-derived extracellular vesicles (MSC-EVs) from eight seminal studies, categorized based on the isolation methods employed: ultracentrifugation-derived EVs (UC-EVs) (Haraszti et al., 2016; La Greca et al., 2018; H. Liu et al., 2019; Salomon et al., 2013), density gradient-derived EVs (HS-EVs) (Bruschi et al., 2018; Haraszti et al., 2016) and a mixed population of EVs (Mix-EVs) (Angulski et al., 2017; Braga et al., 2022; H.-S. Kim et al., 2012). Notably, for Mix-EVs, ultracentrifugation was conducted without prior isolation of HS-EVs.

Mitochondrial protein prediction was conducted across eight proteome reports of EVs utilizing four distinct prediction tools including Target P-2.0 (Armenteros et al., 2019), DeepMito (Savojardo, Bruciaferri, Tartari, Martelli, & Casadio, 2020), DeepLoc 2.0 (Thumuluri, Almagro Armenteros, Johansen, Nielsen, & Winther, 2022), and MitoFates (Fukasawa et al., 2015), in the eight proteome reports of EVs.

### 2.16 MCF-7 treatment and flow cytometry

MCF-7 cells were seeded at a density of 2 × 10^5^ cells per well in 12-well plates and incubated overnight. Subsequently, the wells were treated with equal amounts (20 µg) of HS-EVs, UC-EVs, and mitochondria samples. After the 72-hour incubation period, the cells were detached, followed by centrifugation at 1500 rpm for 5 minutes. The resulting pellet was then washed with PBS. The samples were subjected to staining using the Annexin V/Propidium Iodide (PI) apoptosis assay kit (BioLegend, USA) and subsequently analyzed using a BD FACSCalibur flow cytometer (USA). Data analysis was performed using FlowJo software version 7 (USA).

### 2.17 Statistical analysis

All replicates were true biological replicates. Unless indicated, the statistical significance of differences was evaluated by ordinary one-way ANOVA followed by a multiple comparison analysis with Tukey’s multiple comparison test in SPSS (26.0), and p values <0.05 were considered statistically significant.

## 3 RESULTS

### 3.1 Characterization of MSC-derived EVs through different isolation methods

MSCs were cultured and characterized for surface markers and lineage differentiation as shown in our previous paper (Pakzad et al., 2022). SEM images represented the round morphology of each subpopulation (Figure 1B). DLS was employed to determine the size distribution of each MSC-EV subpopulation, where each subpopulation was heterogeneous but within the normal range for EVs (Figure 1D). The DLS graph revealed that HS-EVs are more heterogeneous than others. We performed western blot analysis to detect the expression of protein markers associated with EVs, including CD9/63/81. (Figure 1C). Quantification of WB bands revealed that the expression of enriched markers of EVs was different among the subpopulations and that the expression of CD9 and CD63 was significantly different among different EVs (Figure 1E).

### 3.2 Three subpopulations of EVs are not exclusively composed of ectosomes or exosomes

Considering albumin (ALB) (mainly derived from serum or platelete lysate supplement) and CANX (derived from endoplasmic reticulum membrane) as the major contamination in conditioned media-EVs (CM-EVs), UC-EVs reproducibly presented the highest amount of EVs specific markers whereas HS-EVs and SU-EVs were contaminated with CANX and ALB, respectively (Figure 2B and Supplementary file 3).

**Figure 2.**
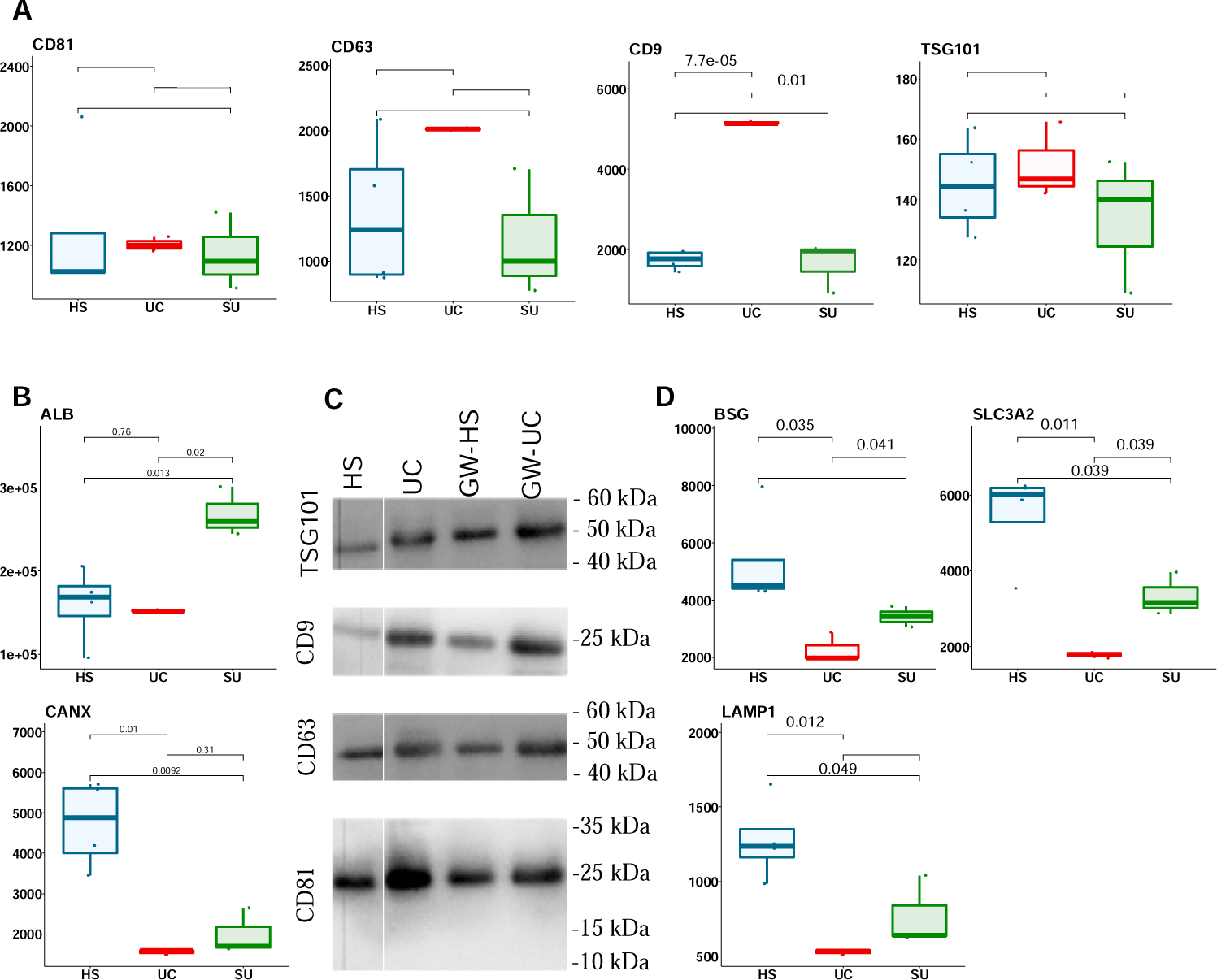
Expression of EV markers in MSC-EV subpopulations identified by mass spectrometry and confirmed via western blotting after inhibiting exosome release. A) Mass spectrometry results for membrane (CD9, CD81, CD63) and luminal (TSG101) characteristic EV markers, and B) negative marker calnexin (CANX). Albumin (ALB) is shown as a major contaminant derived from platelet lysate and showed to be highest in SU group. C) Western blot analysis of specific membrane and luminal protein markers (CD9, CD81, CD63, and TSG101) for HS-EVs (EVs isolated by 20Kg), UC-EVs (EVs isolated by ultracentrifugation on sucrose gradient), treated with and without the specific exosome biogenesis inhibitor GW4869). D) Mass spectrometry-based expression results of LAMP1 as a specific marker of exosome subpopulations, BSG, and SLC3A2 as specific markers of ectosomes. P values were calculated using a t-test in the R platform. HS, EVs isolated by 20Kg; UC, EVs isolated by ultracentrifugation; SU, EVs isolated by ultracentrifugation on sucrose gradient; GW-HS, EVs isolated by 20Kg after MSCs treatment with GW4869; GW-UC, EVs isolated by ultracentrifugation after MSCs treatment with GW4869.

The MS profiles of EVs characteristic CD markers were almost consistent with WB and the amount of TSG101 was very low in all EVs (Figure 2A and Supplementary file 3). Some reports indicated that TSG101 can correspond to the exosome subpopulation of EVs (Hurley, 2010; Tucher et al., 2018; Willms, Johansson, Mäger, Lee, Blomberg, Sadik, Alaarg, Smith, Lehtiö, El Andaloussi, et al., 2016). About EVs biogenesis, exosomes (generated from endosomal pathways) and ectosomes (generated from plasma membrane budding) are the two main subtypes of EVs. Taking into account methodological constraints, particularly the similarities in isolation techniques between UC and SU-EVs, which both utilize ultracentrifugation, we opted to compare UC EVs with HS-EVs. To check out if they are enriched in exosome, we inhibited exosome biogenesis using GW4869, and checked out the expression of all four EV characteristic markers. As shown in Figure 2C, the expression of CD9, CD81, CD63, and TSG101 were the same before and after treatment with GW4869. This may indicate that none of these methods can specifically isolate exosomes. LAMP1 (for exosomes) and BSG and SLC3A2 (for ectosomes) have already been introduced as specific markers for Hela cell-derived EVs (Mathieu et al., 2021). We observed that HS-EVs and UC-EVs contained the highest and lowest amounts of all three markers, respectively (Figure 2D). Totally, it may suggest that these subpopulations of EVs are a heterogeneous subtype composed of exosomes and ectosomes.

### 3.3 Proteome analysis of EVs by mass spectrometry

We used TMT-based mass spectrometry (MS) to quantitatively compare the protein composition of different MSC-EV subpopulations. In total, 2919 proteins were identified and quantified among the EV subpopulations (Supplementary file 1), all were analyzed in biological triplicates. Only the characterized proteins having at least one peptide and FDR <0.01 are presented. The proteomics MS raw data from this study were deposited in the ProteomeXchange Consortium via the GUI-based PX Submission tool under the dataset identifier PXD022174, username: and password: OGfjqI7W.

DAPs between each group were obtained using ANOVA followed by Tukey’s HSD post hoc analysis, and those proteins with a p value <0.05 and log2FC>|0.263| were considered differentially abundant among each group. These analyses revealed 1109, 1504, and 950 DAPs in the HS vs. SU, HS vs. UC, and SU vs. UC comparisons, respectively (Supplementary file 1). The HS vs. UC comparison yielded the greatest number of DAPs, where 1103 proteins were more abundant in HS and 401 proteins were more abundant in UC. Comparison of HS vs. SU found 741 proteins were more abundant in HS and 368 proteins more abundant in SU. The SU vs. UC comparison showed the lowest number of DAPs, where 558 and 392 proteins were highly abundant in SU and UC, respectively. Heatmap plots for each comparison group show the overall consistency among replicates (Figure 3). In addition, there was a significant overlap between DAPs from each comparison group (Figure 3D). However, 73, 41, 264, 49, 109 and 23 proteins showed specifically higher abundances in HS vs. SU, SU vs. HS, HS vs. UC, UC vs. HS, SU vs. UC and UC vs. SU, respectively. Moreever, Principal-component analysis (PCA) of common DAPs between all three EVs group have confirmed the clustering results (Supplementary Figure 1).

**Figure 3.**
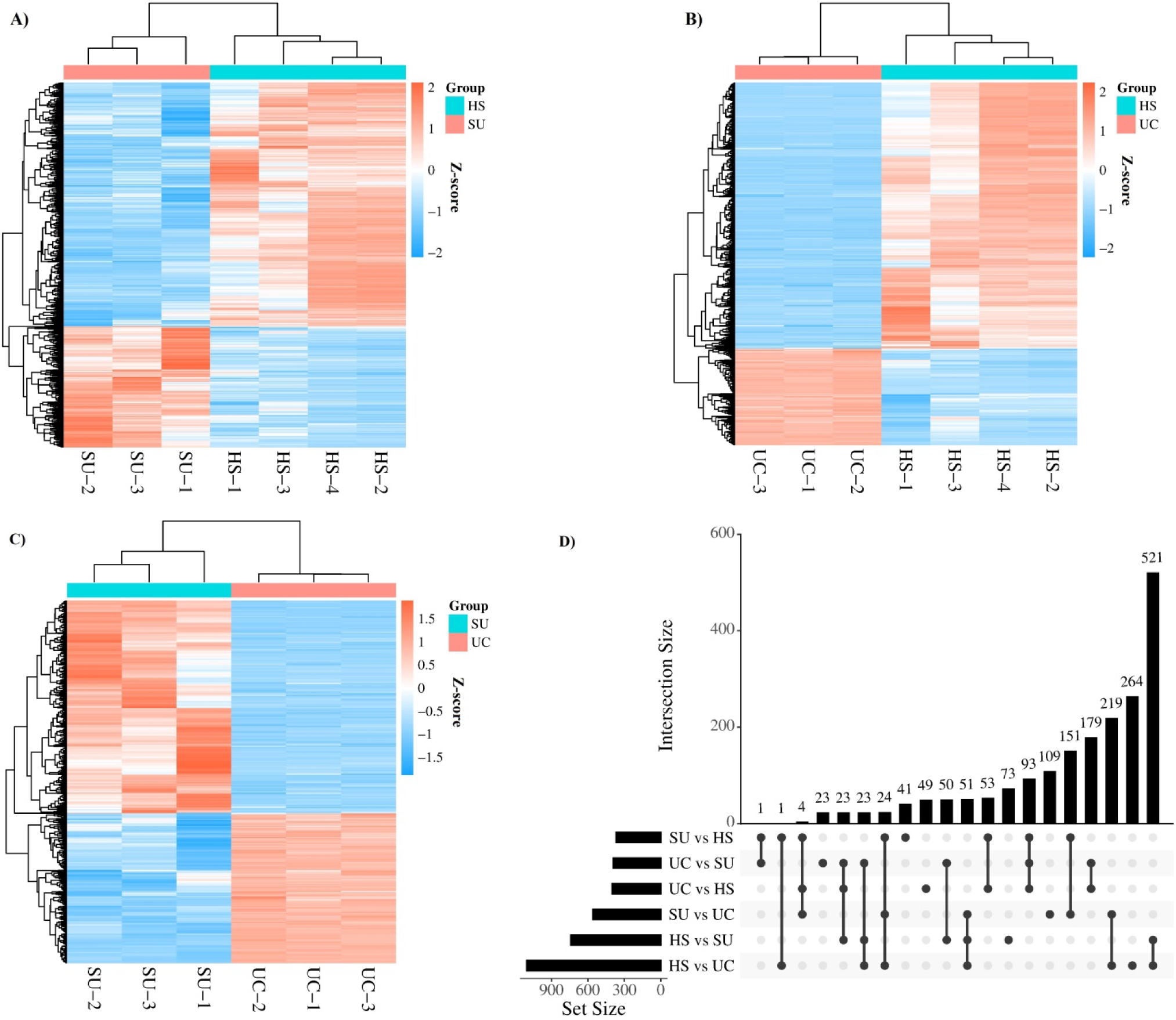
Heatmaps (with hierarchical clustering) A) HS vs. SU, B) HS vs. UC and C) SU vs, UC showing the log-transformed ratios of differentially abundant proteins from three comparison groups, including HS vs. SU, HS vs. UC and SU vs. UC; D) UpSet plot indicating the overlap of proteins between comparison groups.

### 3.4 Functional enrichment analysis revealed key pathways corresponding to DAPs in each comparison group

GO enrichment analysis was performed to identify the biological processes (BP), molecular functions (MF), and cellular components (CC) associated with DAPs in each group. The top three enriched terms are provided in Figure 4(Supplementary file 2). Among the top three terms, a higher abundance of proteins involved in translation and metabolism was observed in the HS group compared to the SU and UC groups (Figure 4 A and C), as well as the SU group compared to the UC group (Figure 4 B). On the other hand, a higher abundance of proteins involved in platelet activation, blood coagulation, and immune response was observed in the UC and SU groups compared to the HS group and in the UC group compared to the SU group (Figure 4 and supplementary file 2). Interestingly, based on the top three identified cellular components, ribosomes were specifically associated with DAPs of EVs from the HS group. On the other hand, the endoplasmic reticulum was observed in the SU group compared to the UC group, and platelets were identified as a common cellular component between the SU and UC groups compared to the HS group.

**Figure 4.**
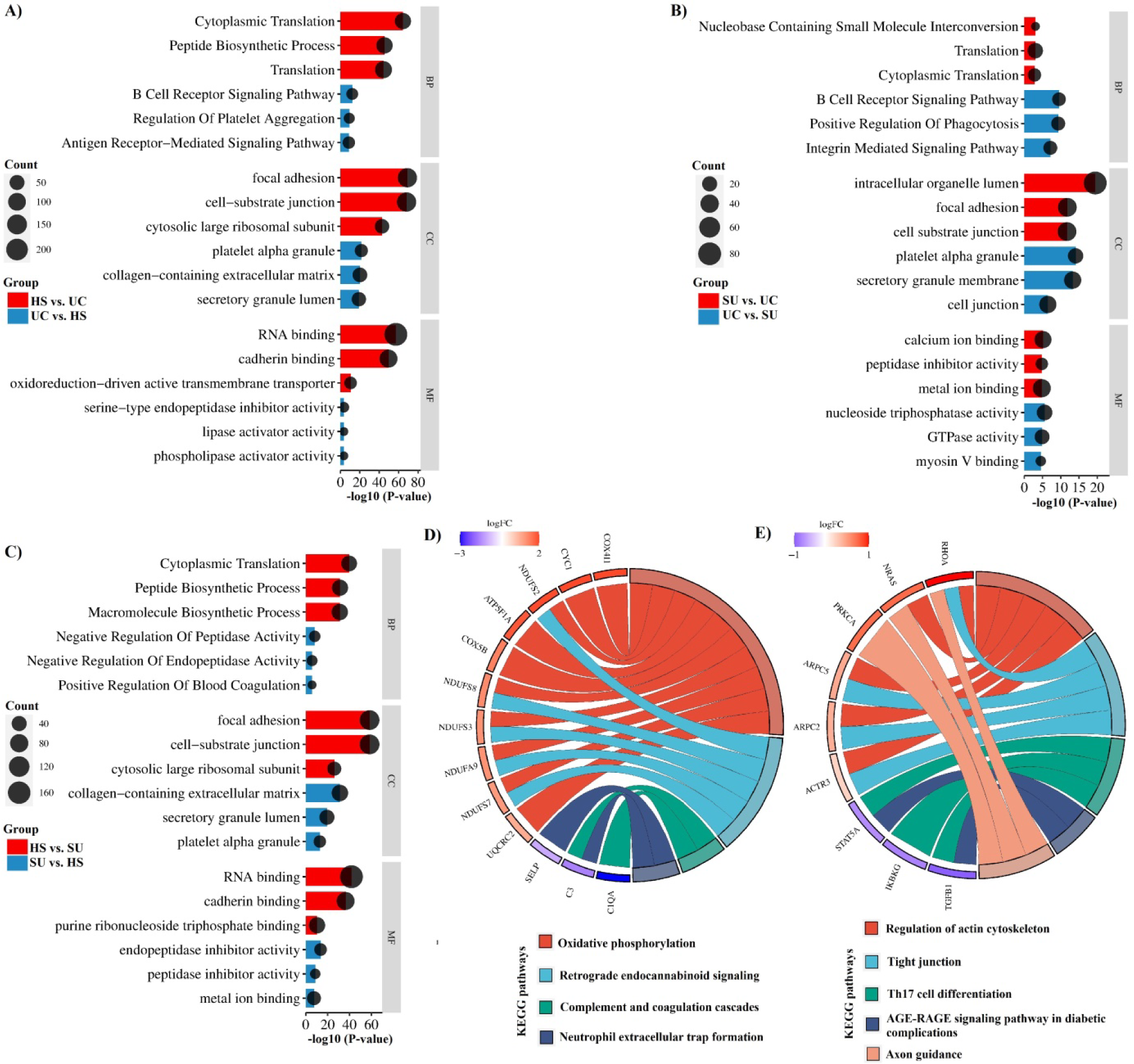
Top 3 most significantly altered biological processes (BP) molecular function (MF) and cellular components (CC) associated with DAPs in A) HS vs. UC and UC vs. HS; B) SU vs. UC and UC vs. SU, and C) HS vs. SU and SU vs. HS; comparison groups. KEGG pathways enriched by the top identified hub genes in the D) HS vs. UC and E) HS vs. SU comparison groups. The oxidative phosphorylation pathway was the main altered pathway, including 10 identified hub genes in the HS vs. UC comparison group.

Additionally, KEGG pathway analysis indicated that the HS group, compared to the SU and UC groups, and the SU group, compared to the UC group, had a higher abundance of proteins involved in translation and metabolism pathways, such as oxidative phosphorylation. On the other hand, the HS groups showed a lower abundance of proteins involved in immune response pathways, such as complement and coagulation cascades, and Th17 cell differentiation. Interestingly, KEGG pathway analysis of the identified hub genes also revealed enrichment in critical pathways for cell homeostasis, including oxidative phosphorylation, retrograde endocannabinoid signaling pathway, regulation of actin cytoskeleton, tight junction, and axon guidance, in the HS group compared to the SU and UC groups. The HS group also showed a lower abundance of proteins involved in inflammatory responses and cell death, such as complement and coagulation cascades, Th17 cell differentiation, the AGE-RAGE signaling pathway in diabetic complications, and neutrophil extracellular traps (NETs) (Figure 4 D and E, and Supplementary file 2).

AGE-RAGE signaling is involved in diabetic complications such as diabetic neuropathy, diabetic nephropathy, and diabetic vascular issues. Moreover, the RAGE signaling pathway plays a significant role in AGE-induced tau phosphorylation and the impairment of spatial memory and has been suggested as a key pathway linking diabetes and AD (Kong et al., 2020). NETs are structures generated by neutrophils as a defensive mechanism to ensnare and neutralize invading pathogens. Nevertheless, these structures can also contribute to the onset of diverse pathophysiological conditions, such as sterile inflammation and autoimmunity (Bonaventura et al., 2018; Papayannopoulos, 2018).

### 3.5 Confirmation of MS result for IGF-1 protein, blood coagulation, and immune response

To confirm the proteomics data, the level of IGF-1, which was found to have different abundances across EV subpopulations, was measured by ELISA. Consistent with the proteomics data, the level of IGF-1 protein was found to be highest in the HS group and lowest in the UC group (Figure 5A). Although our proteomic data and following functional analyses showed that blood coagulation, and immune response were enriched in UC and SU groups compared to the HS group (Figure 4), results of blood coagulation test showed no significant effects of all three EV groups on coagulation process (Figure 5B). However, results of the Lymphocyte proliferation assay (LPA) revealed that different EV types could down-regulate T cell proliferation (Fig 6C). In this regard, SU-EVs significantly exhibited higher anti-proliferative effects.

**Figure 5.**
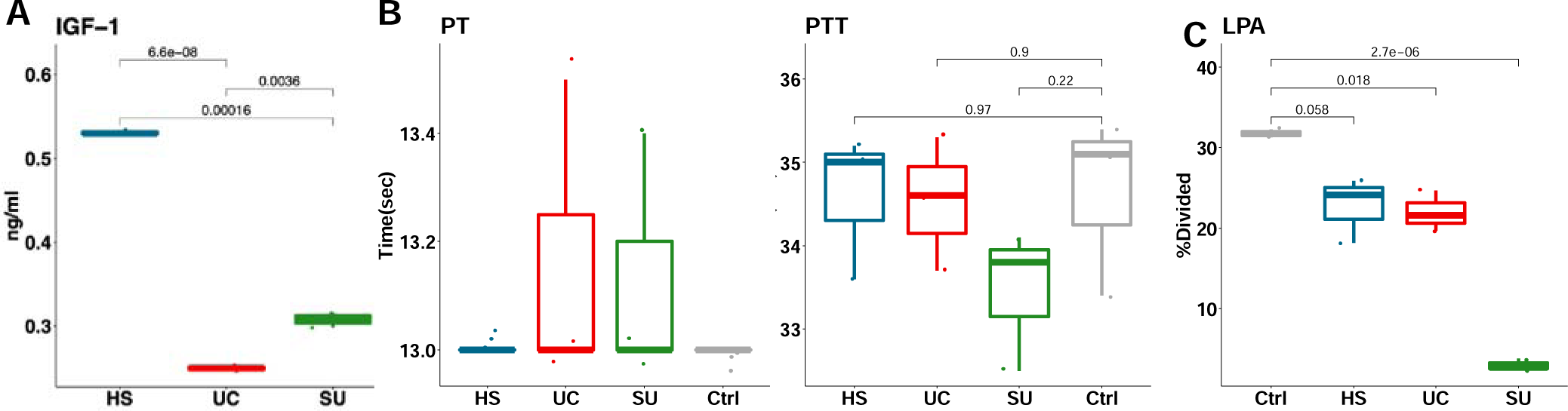
Confirmation of MS results for IGF-1 protein, blood coagulation, and immune response in the EVs subtype. The quantification of IGF-1 (A) is checked in three biological replicates of three MSC-EVs subtypes by ELISA test. (B) Results of PT and aPTT blood coagulation tests on healthy human samples treated with three biological replicates of three MSC-EVs subtypes, compared to Ctrl group that is not treated with EVs. (C) Quantified flow cytometry results of LPA in healthy human T cells treated with three biological replicates of three MSC-EVs subtypes in addition to PHA, compared to Ctrl group that is not treated with EVs and only received PHA. P values were calculated by the t-test in the R platform. HS: EVs isolated by 20Kg, UC: EVs isolated by ultracentrifugation, SU: EVs isolated by ultracentrifugation on sucrose gradient.

**Figure 6.**
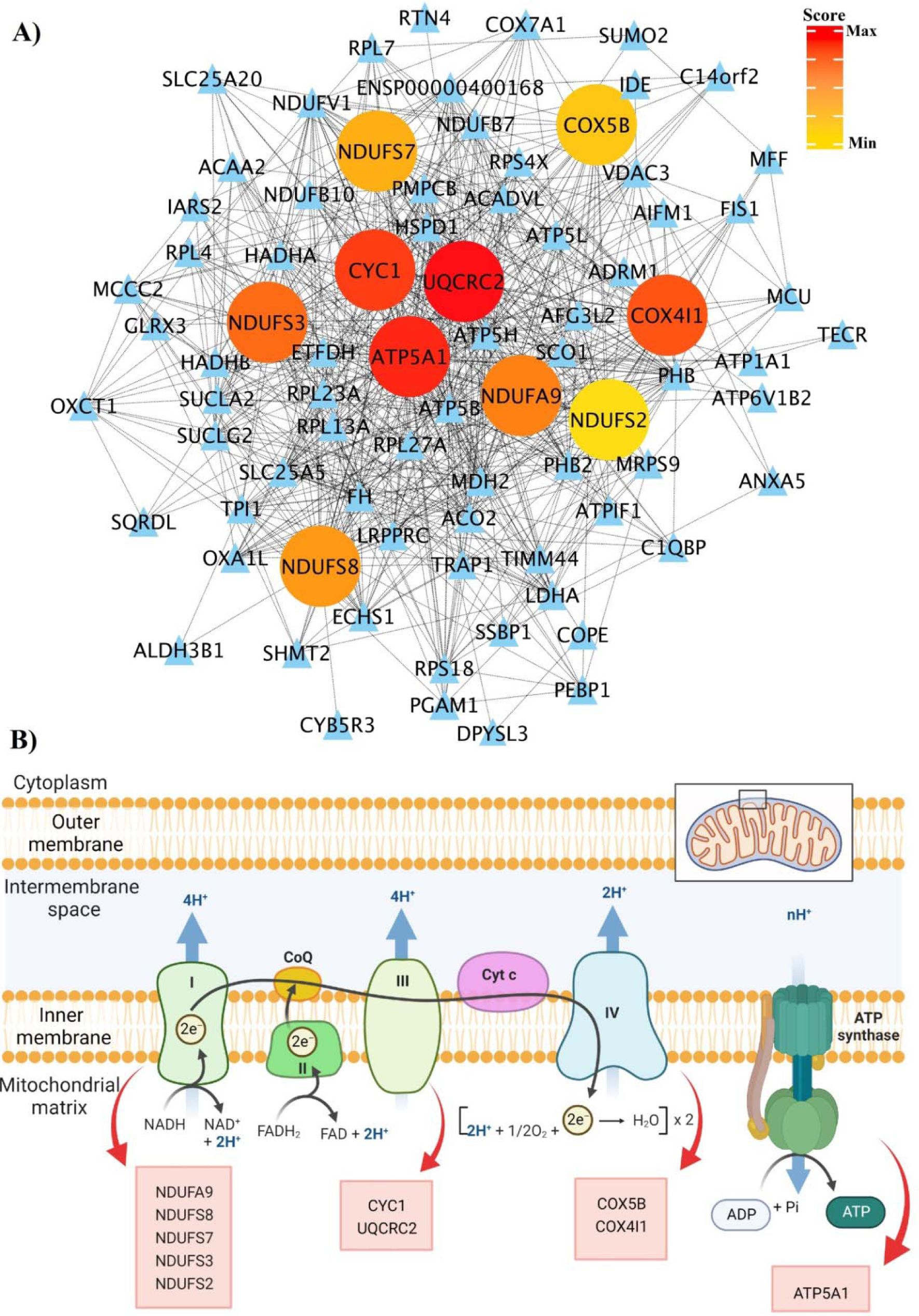
A) PPI network of DAPs in the HS vs. UC comparison group. The top ranked hub genes based on the maximal clique centrality (MCC) algorithm in the cytoHubba plugin of Cytoscape are highlighted from red to yellow based on their resulting score. B) The mitochondrial electron transport chain (ETC) and identified hub genes associated with ETC complexes.

### 3.6 PPI network analysis of DAPs identified hub genes associated with each specific comparison group

To understand the interactions between DAPs specifically in each comparison group and identify hub genes within their networks, we performed PPI interaction and hub gene analysis. Our analyses revealed the top 10 hub genes within each comparison group, as detailed in Supplementary file 1. Among all the identified hub genes, the hub genes within the HS vs. UC PPI network exhibited the highest scores. These hub genes are all mitochondrial proteins related to the electron transport chain, specifically complexes I, III, IV, and ATP synthase (Figure 6). The hub genes include Ubiquinol-Cytochrome C Reductase Core Protein 2 (UQCRC2), ATP synthase F1 subunit alpha (ATP5F1A), cytochrome c1 (CYC1), Cytochrome C Oxidase Subunit 4I1 (COX4I1), NADH:Ubiquinone Oxidoreductase Core Subunit S3 (NDUFS3), NADH:Ubiquinone Oxidoreductase Subunit A9 (NDUFA9), NADH:Ubiquinone Oxidoreductase Core Subunit S8 (NDUFS8), NADH:Ubiquinone Oxidoreductase Core Subunit S7 (NDUFS7), Cytochrome C Oxidase Subunit 5B (COX5B), and NADH:Ubiquinone Oxidoreductase Core Subunit S2 (NDUFS2) (Figure 6 and Supplementary file 3). The PPI network analysis of the UC vs. SU comparison group yielded only four hub genes, which showed the lowest scores. These hub genes are involved in blood clotting and lipid metabolism and include Protein C, Inactivator of Coagulation Factors Va and VIIIa (PROC), Glycoprotein Ib Platelet Subunit Beta (GP1BB), Apolipoprotein A1 (APOA1), and Apolipoprotein F (APOF) (Supplementary file 1).

### 3.7 Meta-analysis of other reports showed HS-EVs enriched in mitochondrial proteins

A total of 4,282 proteins were identified across nine studies, encompassing three Mix-EVs, four UC-EVs, and two HS-EVs (Supplementary Table 1). A total of 675 proteins were identified as potential mitochondrial proteins through the combined use of four prediction tools, namely Target P, DeepMito, MitoFates, and DeepLoc. Remarkably, it was observed that roughly 60 percent of the proteins predicted to be mitochondrial were uniquely identified within HS-EVs (Figure 7C). Mitochondria fraction was free of EVs based on EV characteristic marker assessment by western blotting (Figure 7D). The presence of mitochondria marker protein, ATP5A1, also confirmed the meta-analysis result (Figure 7E), which was sensitive to storage conditions (data not shown).

**Figure 7.**
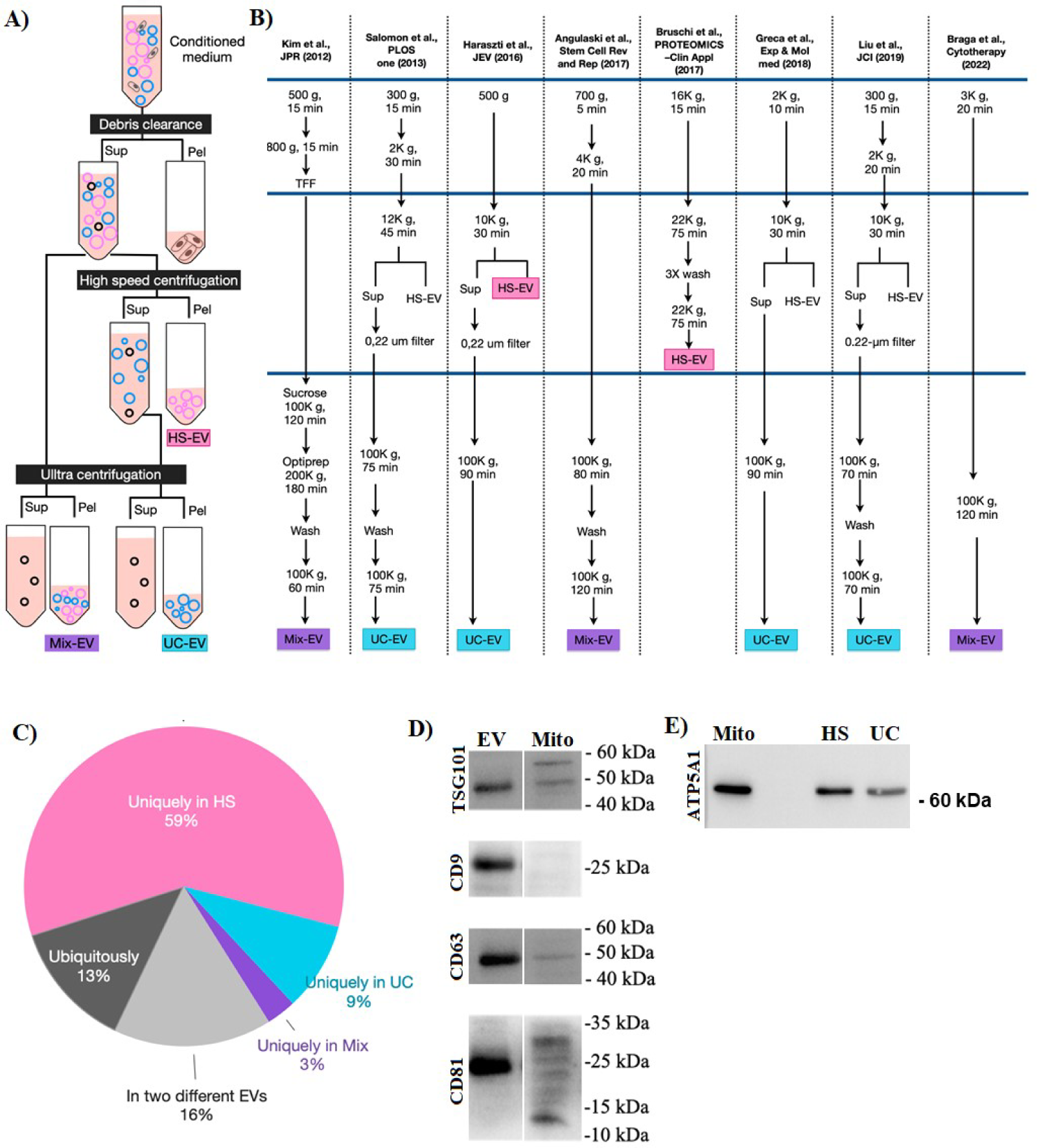
Confirmation of the existence of mitochondrial proteins by meta-analysis followed by mitochondrial prediction using prediction tools and western blotting. A) Overall design of EV isolation by differential centrifugation approach. The first step is clearance of cellular debris by low-speed centrifugation and the second step is high-speed (HS) centrifugation. Finally, UC (with or without density gradient) may isolate the remained EVs. B) Details of EV isolation process in reports with available proteome profile. Pink and blue boxes are proteome reports on HS and UC respectively and purple boxes showed the mixed population of HS and UC separated EVs. C) Various prediction tools were employed to forecast proteins associated with mitochondria including Target P, DeepMito, MitoFates, and DeepLoc. D) The presence of EV characteristic markers in Mitochondria sample compared to EVs by western blotting. E) The expression of ATP5A1 in HS-EVs and UC-EVs. Mitochondria was loaded as positive control. The whole blot image is presented as supplementary Figure 2. HS, EVs isolated by 20Kg; UC, EVs isolated by ultracentrifugation; SU, EVs isolated by ultracentrifugation on sucrose gradient; Mito, mitochondria fraction isolated from MSCs.

### 3.8 HS-EVs but not other EVs effectively induced early apoptosis in cancer cells

Since mitochondrial proteins were found to be key components of EV subpopulations, particularly HS-EVs, the potential cytotoxicity of different EV subpopulations was compared with isolated mitochondria. Breast cancer cell lines were treated with EV subpopulations and mitochondria, and their potential to induce apoptosis was investigated using flow cytometry. As depicted in Figure 8, mitochondria significantly induced both early and late apoptosis in cancer cells. In contrast, cells treated with HS-EVs and UC-EVs did not show significant late apoptosis. However, HS-EVs demonstrated considerable efficacy in inducing early apoptosis (Figure 8). All groups showed a maximum of about 10% necrosis, which was expected (Figure 8).

**Figure 8.**
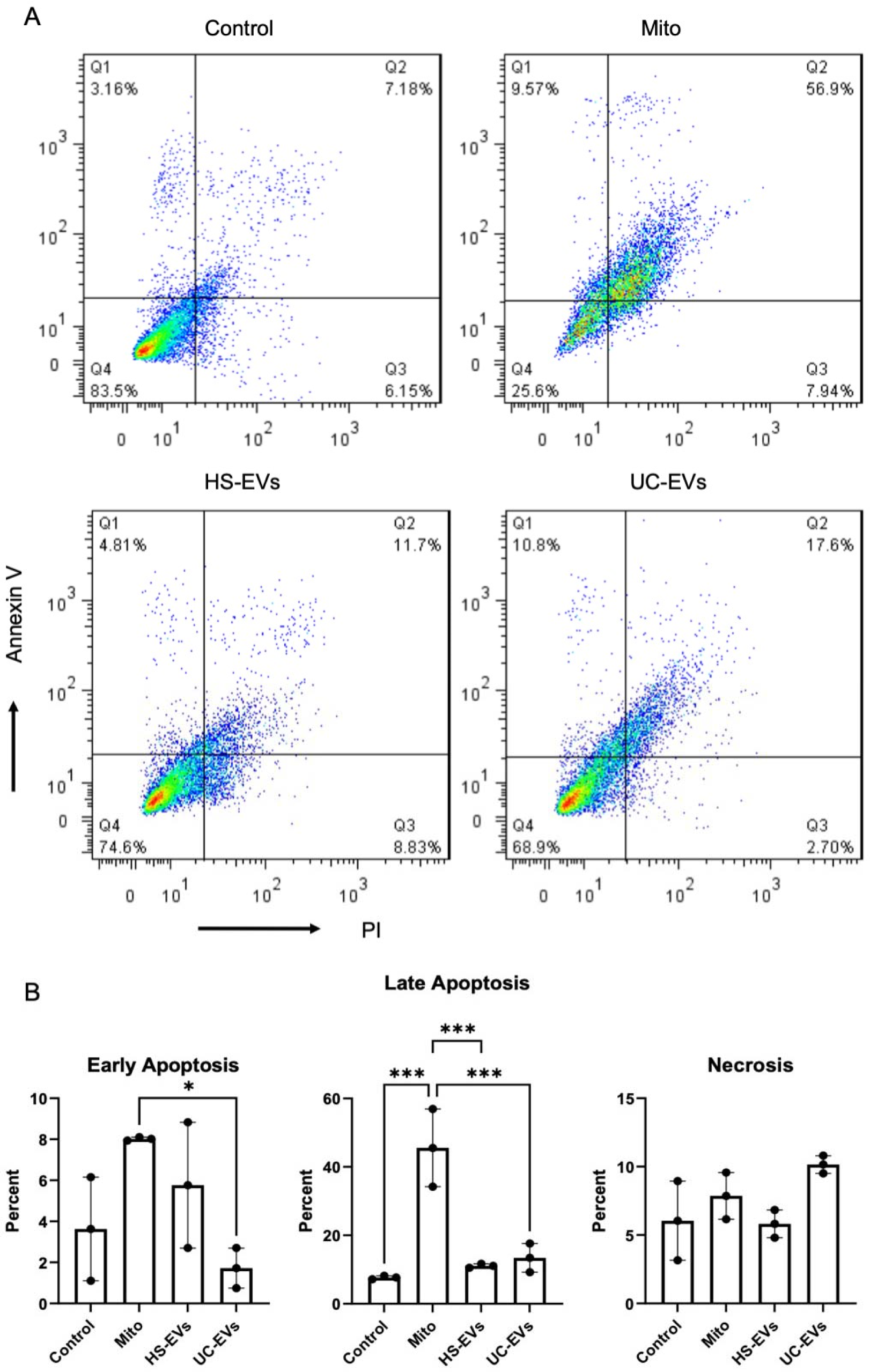
Functional characterization of EVs compared to isolated mitochondria fraction, A) The flow cytometry result of cytotoxicity profile of MCF7 cell line treated with HS-EVs, UC-EVs, and Mitochondria for 24h assessed by annexin/PI test. The four distinct quadrants (Q1, Q2, Q3, and Q4) represent necrosis, late apoptosis, early apoptosis, and live cells, respectively; B) Bar graph illustrating the percentages of early apoptosis, late apoptosis, and necrosis in each group. Data are presented as mean percentages ± standard deviation (SD).

## 4 DISCUSSION

In this proteomics study, we conducted a comprehensive quantitative proteomic analysis of MSC-derived EVs obtained using commonly employed isolation methods. Our findings revealed substantial variations in the proteome contents of EVs isolated through different methods, thus emphasizing the potential implications of these variances for therapeutic applications. We acknowledged the multifaceted nature of EVs and their diverse cargo, recognizing that EV proteomics alone may not fully capture the complexity of their therapeutic potential. However, this proteome report may provide insights into the differential protein content of EVs based on the isolation method.

Each cell releases diverse subpopulations of EVs that are distinct in size, content, and function (Kalluri & LeBleu, 2020; Kowal et al., 2016; Willms, Johansson, Mäger, Lee, Blomberg, Sadik, Alaarg, Smith, Lehtiö, & El Andaloussi, 2016). MSC-EVs have been exploited for prominent therapeutic benefits in regenerative medicine and have been used in clinical trials (Kordelas et al., 2014; Kwon et al., 2020; Nassar et al., 2016; K. Y. Park et al., 2021; Sengupta et al., 2020) and preclinical studies (reviewed in (Akyurekli et al., 2015; Elahi, Farwell, Nolta, & Anderson, 2020; F. Shekari et al., 2021)). However, as highly recommended by ISCT and ISEV, more critical investigations are required before administration in a clinical trial (Börger et al., 2020). As shown in Figure 1, UC, HS and SU EVs are three distinct subpopulations since their MISEV2018 characteristics, such as size and marker expression, differ. The UC-based EV isolation seems to be the most reproducible, as shown by three replicates of western blotting. Recently, it has been shown that small EVs bearing tetraspanins, especially CD9 and CD81 with little CD63, originate mainly from the plasma membrane, whereas others bearing CD63 with little CD9 are qualified to be named exosomes (Mathieu et al., 2021). All three markers were present in HS, UC and SU EVs, while for western blotting (Figure 1), the highest level belonged to UC-EVs. Therefore, it seems that these three subpopulations are a mix of ectosomes and exosomes.

We selected UC as the most routinely used method, SU as the method with the low pressure on EVs, and HS as the feasible method that can be performed more easily in clinical GMP grade settings. Based on western blotting, size distribution (Figure 1), and MS results (Figure 2), the UC-based EV isolation method seems to be the most reproducible one, with the lowest albumin and calnexin contamination (Figure 2).

The functional enrichment analyses of all identified proteins and hub genes within the PPI network in each comparison group revealed certain patterns. Specifically, the HS group showed a higher abundance of ribosomal proteins and proteins involved in translation and metabolism compared to the SU group, and both the HS and SU groups had higher abundance in these categories compared to the UC group. Additionally, the HS group exhibited a higher abundance of proteins involved in oxidative phosphorylation and retrograde endocannabinoid signaling (which shares most of the proteins with oxidative phosphorylation pathway) than both the SU and UC groups. On the other hand, the results indicated a lower abundance of proteins involved in immune response pathways, such as complement and coagulation cascades, Th17 cell differentiation, the AGE-RAGE signaling pathway in diabetic complications, and NETs in the HS group compared to both the SU and UC groups, as well as in the SU group compared to the UC group. Our results were in line with the literature that indicated a higher abundance of ribosomal and mitochondrial proteins in large EVs than in small EVs (Jimenez et al., 2019; Lischnig et al., 2022). Moreover, we verified the presence of mitochondrial proteins in EVs through both meta-analysis and in vitro experiments (Figures 8 and 9). To our knowledge, this study is the first to compare mitochondrial content across different EV subpopulations. These preliminary findings pave the way for the potential use of EVs derived from mesenchymal stem cells in conditions such as metabolic disorders and cancer.Our results also revealed a higher abundance of immune-related proteins in EVs isolated through the UC and SU methods than in those isolated through the HS method. We validated the immunomodulatory effects of EVs isolated through HS, UC, and SU using the LPA assay. LPA, a widely employed method for assessing cell-mediated immune responses, has previously demonstrated the immunomodulatory potential of both MSCs and their EVs (Pachler et al., 2017). Accordingly, the immunomodulatory effect of different EV subtypes was investigated by this test and it was observed that different EV subtypes effectively suppress the lymphocyte proliferation as an immune response criterion (Figure 5C). The complement system (CS), an integral component of the human immune system, plays a crucial role in defending against pathogens. Additionally, emerging evidence suggests the involvement of the complement and coagulation cascades in various diseases, including infectious and neurodegenerative conditions (Ma et al., 2021). Notably, complement C3, which is a key player in the complement cascade, exhibited significantly higher abundance in the UC group than in the HS group. Complement C3 mediates synapse elimination in the central nervous system (CNS) and brain aging, since CS is involved in synaptic refinement during brain development through the removal of weak synapses (Shi et al., 2015). A significantly higher abundance of complement C6, which has a key role in membrane attack complex (MAC) assembly, was observed in the UC group than in the HS group. MAC was shown to directly damage synapses in an Alzheimer’s disease (AD) model, and C6 knockout was associated with significantly reduced synapse loss (Carpanini et al., 2022). The presence of complement factors in EVs that isolated by high speed centrifugation (irrespective to the name that used in the literature) has been reported previously (Biró et al., 2007; Karasu, Eisenhardt, Harant, & Huber-Lang, 2018; Kilpinen et al., 2013).

The NET formation pathway was another enriched pathway in the UC group compared to the HS and SU groups. Neutrophils play a key role in pathogen clearance through phagocytosis, NETs, and reactive oxygen species (ROS) generation (Vitte, Michel, Bongrand, & Gastaut, 2004). Increased numbers of neutrophils and proteins associated with neutrophil degranulation have been widely reported in infectious diseases such as SARS-CoV-2 (Akgun et al., 2020; Rosa et al., 2021). Neutrophil degranulation has been identified as one of the main enriched pathways in proteomics analysis of COVID-19 patients (Bankar et al., 2021). EVs derived from MSCs have been demonstrated to possess antimicrobial effects and can effectively enhance the clearance of microbes in an animal model of pneumonia (Xiaolan Wu et al., 2022). However, the specific molecular mechanisms underlying these effects are not yet fully understood. One potential mechanism that contributes to the antimicrobial effects of MSC-derived EVs may be the induction of NET formation. Intriguingly, a recent study showed that NETs and their histones directly promote Th17 cell differentiation, which is another enriched pathway in SU compared to HS groups that also plays a key role in protection against infections at mucosal surfaces (Khader, Gaffen, & Kolls, 2009; Wilson et al., 2022). A significantly higher abundance of STAT3, which is a critical factor for Th17 differentiation, was observed in the SU group than in the HS group. Although Th17 cells are well known as proinflammatory cells and aberrant activation has been linked to the pathogenesis of most common autoimmune diseases, such as multiple sclerosis (MS) (Huang, Wang, & Chi, 2012), they can be potential anti-inflammatory cells depending on their environmental changes (Xu et al., 2020).

Enrichment of the AGE-RAGE signaling pathway in diabetic complications is another significant finding of this study, which was observed in both the SU and UC groups compared to the HS group. The AGE–RAGE signaling pathway is mainly known for its role in diabetic complications, including diabetic osteopathy; however, its role in other diseases, such as AD, cardiovascular diseases (CVDs) and cancer, has also been reported (Rungratanawanich, Qu, Wang, Essa, & Song, 2021). On the other hand, enrichment of oxidative phosphorylation was observed in the HS group compared to both the SU and UC groups. This finding is consistent with previous reports indicating enrichment of mitochondria in large EVs obtained through 10,000 g centrifugation followed by filtration (Ikeda et al., 2021). The term “mitovesicles” refers to extracellular vesicles (EVs) that carry mitochondrial components, such as proteins, lipids, and mitochondrial DNA (mtDNA) (D’Acunzo et al., 2021). These EVs exhibit unique characteristics in terms of morphology, as well as protein and lipid composition, distinguishing them from microvesicles and exosomes (D’Acunzo et al., 2022). While evidence of mitochondrial content has been previously reported in other cell types, (Rai, Fang, Claridge, Simpson, & Greening, 2021; Wang et al., 2020), our study conducted a comparative analysis of different MSC-EVs subpopulations. Our findings revealed that mitochondrial proteins were present in all EV groups, with a higher abundance observed in the HS group compared to the other groups. Oxidative phosphorylation is an important metabolic pathway and a major source of energy that occurs in the mitochondria. Mitochondrial dysfunction, leading to the deregulation of the OXPHOS pathway, has been identified as a key pathway affected in various diseases, such as AD, CVDs, Parkinson’s disease, MS, and diabetes (Abyadeh, Gupta, Chitranshi, et al., 2021). Consequently, there has been a significant focus on intervention approaches aimed at targeting mitochondria as potential therapeutic options for these debilitating diseases. Notably, the beneficial effects of EVs derived from MSCs in improving mitochondrial function have made EVs an intriguing target in this regard (Park, Shin, Duong, Lee, & Lew, 2022; Silva et al., 2021; Zhao et al., 2020). Comparing the beneficial effects of mitovesicles, which specifically carry mitochondrial proteins, with various EV subpopulations that not only contain several mitochondrial proteins but also include proteins related to the anti-inflammatory pathway, is indeed interesting for future studies. However, based on our results, mitochondria and mitovesicles may be a better option for addressing cancer, while EVs are probably a better option for neurodegenerative diseases like AD. Moreover, retrograde endocannabinoid signaling, another affected pathway in AD, has been observed to be more prevalent in HS compared to other EV subpopulations. Endocannabinoids play a significant neuroprotective role in the AD model by regulating the timing of excitatory and inhibitory synaptic neurotransmitter release. They bind to and stimulate the cannabinoid 1 receptor (CB1R), thereby promoting synaptic plasticity and efficient neurotransmitter transmission, crucial for cognitive functions, memory, emotions, and motor skills (Abyadeh, Gupta, Paulo, et al., 2021; F. Chen et al., 2022). It’s important to note that mitochondrial transfer/transplantation has implications for tumorigenesis, drug resistance, and chemotherapy efficacy in cancer treatment. Additionally, mitochondria play crucial roles in various cell death mechanisms, acting as both sensors and amplifiers. When mitochondria become denser, there’s a risk of increased harmful reactive oxygen species (ROS) production and higher levels of cytochrome C, potentially hindering cell clearance and promoting apoptosis. Conversely, research indicates that acquired mitochondria can elevate endogenous mitochondrial DNA levels. Excess mitochondria may lead to surplus ATP production, surpassing the organism’s energy requirements and exerting adverse effects (Z. Liu, Sun, Qi, Cao, & Ding, 2022).

In conclusion, the results of the present study provide insights into the proteome content of MSC-derived EVs isolated through different methods. These findings can be instrumental in guiding the selection of EV subpopulations based on the desired features. It is noteworthy that our results revealed a higher abundance of mitochondrial and ribosomal proteins in the HS group than in the SU and UC groups, while inflammatory proteins showed a lower abundance. Our results suggest that EVs isolated using the HS method with a wider size distribution, including both small and large EVs, may show better therapeutic features than EVs isolated through other methods. It is important to acknowledge that the majority of proteins were shared among all three groups, with differences primarily observed in their abundance. Therefore, the application of EVs isolated through either high-speed centrifugation or ultracentrifugation may exhibit therapeutic effects, given that both small and large EVs have shown therapeutic potential in existing literature. (Dave et al., 2023; Ikeda et al., 2021; Park et al., 2022; Silva et al., 2021). Therefore, considering the proteome content of EV subpopulations could enhance their therapeutic application.

In this investigation, we employed clonal MSCs as a uniform source, recognizing that our observations might be specifically relevant to MSCs-derived EVs. To broaden the applicability of these findings to EVs from various cell types, additional investigations incorporating diverse cell sources are essential. Furthermore, our testing of EV subpopulations was limited to cancer cells, serving as a proof of concept for the functionality of EVs. Subsequent studies are needed to ascertain whether these outcomes can be reproduced in other cell lines and EVs derived from different origins. On the other hand, as emphasized in the MISEV2023 guideline, the molecular and structural heterogeneity of EVs presents challenges at every stage of EV studies, especially in the context of therapeutic or diagnostic applications (Welsh et al., 2024). Considering the multitude of parameters from cell culturing that influence this heterogeneity, as discussed recently, this issue is becoming increasingly complex (Faezeh Shekari, Faisal J. Alibhai, et al., 2023). Previous discussions have suggested that employing single-vesicle technologies can unveil the heterogeneity of EVs (Ferguson, Yang, & Weissleder, 2022; Ferguson, Yang, Zelga, et al., 2022; Sun et al., 2023). Perhaps, conducting large-scale proteome profiling of EVs at the single EV level could offer more robust evidence in this regard.

Overall, our data lay the foundation for further analysis of the therapeutic and diagnostic applications of EVs. However, additional studies exploring other components of EVs, such as miRNAs, are warranted to unravel the secrets within these membrane-bound vesicles and pave the way for their application in clinical settings.

## Supporting information

In total, 2919 proteins were identified and quantified among the EV subpopulations (Supplementary file 1)

The top three enriched terms are provided in Figure 4(Supplementary file 2)

## ACKNOWLEDGEMENTS

The authors have nothing to report.

---

**Supplementary Figure 1.**
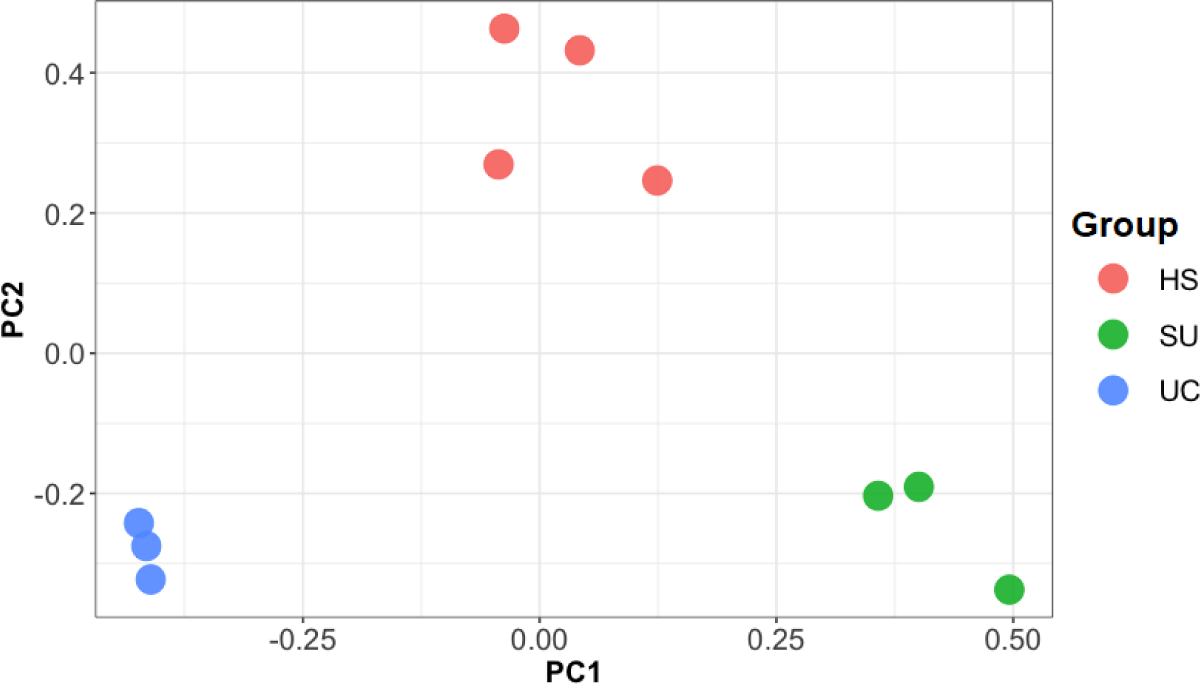
Principal-component analysis (PCA) of common differentially abundant proteins between all three experimental groups, including HS, SU and UC.

**Supplementary Table 1:**
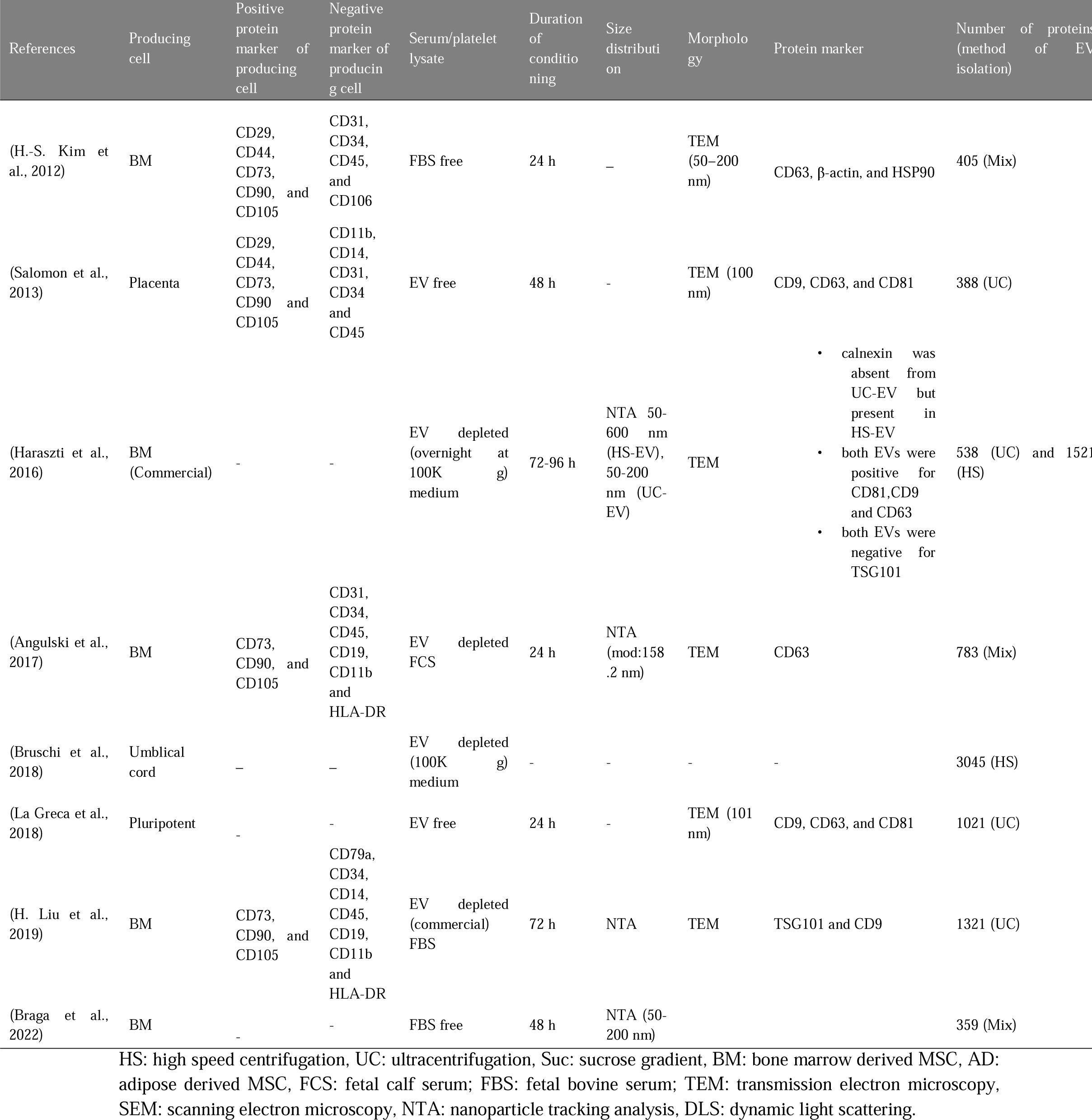
Proteome profiling reports of centrifugation based MSC-EVs.

